# Variation in diel timing in great tits is affected by the timing of their social mate

**DOI:** 10.1101/2024.08.26.609680

**Authors:** Aurelia F. T. Strauß, Barbara M. Tomotani, Marcel E. Visser, Barbara Helm

## Abstract

Diel rhythms are driven by genetic and environmental components. These rhythms are mediated by the circadian clock, and entail rhythmicity of various physiological and behavioural traits. Although individuals show to some extent repeatable timing (i.e., chronotype), there is ample variation of diel timing observed within and between individuals of the same species. Here, we investigated various environmental factors, including timing of the social partner, that could explain day-to-day variation within individuals. Synchronisation with the social partner during provisioning timing could increase breeding success, and decrease extrapair paternity opportunities for females during the fertile period in case of consistent timing across the breeding stages. Therefore, we also investigated fitness consequences of between-individual variation. We first assessed the magnitude of between- and within-individual variation in the timing of nest visits by great tits (*Parus major*). We monitored nest visits of males and females in 37 broods during chick provisioning in 2020 and 2021. Next, we explored the responsiveness of the diel timing to environmental variables, specifically comparing abiotic and social factors. The onset of nest visits varied significantly with day within the breeding season, rainfall and the diel timing of the breeding partner but not with night temperature. In response to the partner’s onset, females responded stronger compared to males. By contrast, offset was generally more variable within individuals and less variation was explained by the environmental variables. Both males and females delayed their activity offset with the progressing season and females also had a later onset with more daytime rainfall. Further, the reproductive output and extrapair paternity were independent of parental chronotype and their synchronisation within pairs. It is possible that consistency of chronotypes is less important for reproductive success than the ability to plastically respond to changing environmental conditions. Thus, the next step could be to investigate potential individual differences in plasticity which could be even linked to specific chronotypes. This information might be crucial to predict how species can cope with unpredictable environmental conditions.

## Introduction

Activity patterns show rhythmic repetition every 24 h (= diel) across the animal kingdom (Anderson & Wiens, 2017). These diel rhythms are driven by the internal circadian clock that sustains rhythmicity under constant conditions and entrains to a 24-h rhythm using environmental cues, predominately day-night transitions (Aschoff, 1967; Cassone, 2014). The genetic background determines the speed (i.e. period length) of the circadian clock (Cassone, 2014; Steinmeyer et al., 2012 but see Laine et al., 2019) as well as other mechanisms that are involved in the long- and short-term adjustment of the rhythm to the environment (Helm et al., 2017). There is ample variation in diel timing not just between species, e.g. nocturnal owls vs. diurnal songbirds, but also between individuals of the same species. Consistent between-individual differences in diel timing of activity are commonly referred to as chronotypes, usually varying from early to late (Schwartz et al., 2017). These differences have been found to account for up to 46% of the variation in diel timing in birds (Lehmann et al., 2012; Maury et al., 2020; Meijdam et al., 2022 but see Strauß et al., pending revision; Stuber et al., 2015). Additionally, there is variation within individuals that plastically adjust their diel timing on a day-to-day basis (Helm et al., 2017). For example, activity patterns can adjust to external cues such as weather conditions (Schlicht & Kempenaers, 2020) or conspecifics (Favreau et al., 2009).

While seasonal changes in diel timing, as well as plastic responses to weather conditions, are known (McGlade et al., 2023; Schlicht & Kempenaers, 2020; Strauß et al., 2024), the effects of conspecifics on the circadian clock and diel timing are less studied, in particular in wild animals (Bloch et al., 2013; Davidson & Menaker, 2003; Favreau et al., 2009; Regal & Connolly, 1980). If other cues are absent, the internal clock can entrain to social cues (Gwinner, 1966; Menaker & Eskin, 1966; Regal & Connolly, 1980; Shimmura et al., 2015; Steiger et al., 2013). Social cues can synchronise rhythms and timing of individuals depending on their relationship to each other (Bloch et al., 2013; Favreau et al., 2009). For example, subdominant individuals often desynchronise with dominant individuals to avoid conflicts (Meerlo et al., 2002; Regal & Connolly, 1980).

In the wild, bird studies have focused on synchronisation of parental care which was mostly studied in the alternation of visits by the two parents. For example, in many songbirds both parents take turns in chick provisioning synchronising their nest visits to decrease predation risk and monitor their partner’s activity (Bebbington & Hatchwell, 2016; Johnstone et al., 2014; Lejeune et al., 2019; Leniowski & Wȩgrzyn, 2018; van Rooij & Griffith, 2013). Another example is the turn-taking of the clutch incubation in the lesser yellowleg, *Tringa flavipes*, where females incubate during daytime and males at night (Bulla et al., 2016). Only some studies have investigated the synchronisation of diel timing such as the awakening time of blue tits, *Cyanistes caeruleus*, in winter (Steinmeyer et al., 2013), or the first nest visit per day of great tits, *Parus major*, during provisioning (Pagani-Núñez & Senar, 2016). In both studies, the timing of the breeding pair was positively correlated in which a later onset was associated with a later onset of the partner. However, it is unclear whether this is due to active adjustment towards the partner’s timing, to assortative mating, or to the shared environment of the breeding pair. This is a common issue when correlating phenotypes of breeding pairs (Holtmann & Dingemanse, 2022).

While diel timing is generally considered adaptive, synchronisation in diel timing between social monogamous pairs could also be beneficial (Duranton & Gaunet, 2016; Hau et al., 2017). There is evidence that great tits with an early chronotype raise more offspring (Womack et al., 2023 but see Strauß et al., pending revision.) which could be associated with a higher feeding rate of early-active parents (Pagani-Núñez & Senar, 2016). Early-active blue tit males also gain more extrapair paternity (EPP) during the mating phase in the early morning (Poesel et al., 2006; Schlicht et al., 2023), while experimentally delayed great tit males lost paternity in their own nest (Greives et al., 2015). These late males probably lost the opportunity for mate guarding that reduces a male’s risk that their female mates with other males (Kluijver, 1950). Nevertheless, EPP levels did not increase for experimentally advanced male and female blue tits (Santema & Kempenaers, 2023; Schlicht et al., 2014). This underlines the importance of synchronised timing between male and female in a breeding pair. However, the direct fitness effects of synchronisation of the parental chronotypes have not been studied yet (but see Steinmeyer et al., 2013).

In this study we aim to investigate the sources of variation in diel rhythms of great tits during chick provisioning and their fitness consequences. Therefore, across two breeding seasons (2020-2021) we extracted the diel timing of parents whose visits at the nest-box were monitored. From this data, we first assessed how much variation of the diel timing was explained by between- and within-individual variation using the pair as approximation for the individual (i.e. between- and within-individual variation within pair; see ‘*Variation partitioning*’). We expect repeatable individual differences, with potential differences in the estimates depending on sex as well as between onset and offset of provisioning (Kluijver, 1950; Lehmann et al., 2012; Stuber et al., 2015).

Thereafter, we compared the effects of different environmental factors on diel timing with special attention to the effect of the social partner (see ‘*Environmental factors*’). We assumed that individuals respond differently in their onset and offset to the progressing season and weather conditions, such as ambient temperature and rainfall, and we furthermore anticipated sex differences in responsiveness (Schlicht & Kempenaers, 2020; Stuber et al., 2015). Environmental effects on diel timing can represent between-individual differences in the environment as well as within-individual modification towards the environment. Thus, we compared the repeatabilites of the model with and without environmental effects to confirm that potential effects represent responses within individuals. As for the effect of the social partner, a strong correlation of the breeding partners’ onsets of provisioning has been found previously (Pagani-Núñez & Senar, 2016). However, to disentangle potential confounding factors such as the shared environment, we strived to test for a partner effect while accounting for a set of selected environmental factors (Holtmann & Dingemanse, 2022). If the correlation is a by-product of the shared environment, we expected a reduced correlation between partners.

Lastly, we investigated fitness consequences, i.e. differences in the reproductive output, for male and female chronotypes as well as for the matching of the breeding partners (see ‘*Reproductive output*’). For this analysis, chronotypes were calculated from the onsets of provisioning only because they have been previously shown to represent between-individual differences better than offsets (Stuber et al., 2015) and seem more important for the mate attraction in the morning (Mace, 1989). If an earlier chronotype was beneficial for the reproductive output (Pagani-Núñez & Senar, 2016; Womack et al., 2023), we expected a higher number of fledglings for those chronotypes. Alternatively, desynchronisation of the breeding partners could be beneficial due to an increased time period of paternal availability per day for chick provisioning (Steinmeyer et al., 2013). Under the assumption that chronotypes are consistent throughout the whole breeding season, there might be additional effects on the EPP levels in such pairs. As the relationship of chronotype and EPP levels could be sex-dependent (Schlicht et al., 2023 vs. Schlicht et al., 2014), variation in EPP levels might be explained by the matching of parental chronotypes. A synchronised timing might be crucial for mate guarding during the female’s fertile period (Kluijver, 1950) resulting in lower EPP levels. In turn, desynchronisation would allow the male and the female to have extrapair copulations, increasing EPP levels.

## Methods

### Study area and brood monitoring

We recorded nest visits of 37 broods of great tits (*Parus major*, Linnaeus 1758) from a population on Vlieland (53° 18′ 0″ N, 5° 4′ 0″ E), the Netherlands, in spring 2020 and 2021. This population has been monitored from 1955 (Kluyver, 1970) at least once a week across all breeding stages (i.e. from nest building to fledging). In particular, we recorded lay date (i.e. day of the first egg back-calculated assuming one egg laid per day), hatch date, clutch size and the number of hatchlings and fledglings. On chick age 7-11 days (with hatch date being chick age 0), adults were caught using spring-loaded traps, individually ringed, and their biometrics and a blood sample were taken. For this study, individuals additionally received a passive integrated transponder (PIT) tag (2.6-mm EM4102 PIT bird tag, Eccel Technology Ltd., Leicester, UK), attached to the leg, to record nest visits using a radio-frequency identification (RFID) system (LID665/650 stationary decoder, Dorset ID, Aalten, the Netherlands). Seven females had previously received a PIT tag when they were caught during incubation day 8 using a nest-box net (only in 2021; te Marvelde et al., 2011). We placed the RFID antenna at the nest-box entrance either one day after catching or as soon as we observed an adult with PIT tag from the previous year. We recorded nest visits for both parents in 30 of the 37 broods of which one pair was recorded in both years (n = 29 pairs). Further, six individuals had a different partner in the two years so we measured 26 males and 33 females in total. The chicks were caught for individually ringing (chick age 7-12 days) and to record their biometrics when chicks were 14-16 days old (Verboven & Visser, 1998). Additionally, we took blood samples from adults of 25 broods and from 109 chicks of 23 broods to assess extrapair paternity. One brood was excluded due to insufficient information about the social father and, thus, the final sample size was 105 chicks of 22 pairs.

### Extrapair paternity analysis

Extrapair paternity was assessed using the blood samples taken from adults and their chicks following a standard protocol (Greives et al., 2015; Helm & Visser, 2010). In specific, we isolated the genomic DNA from the blood samples (FAVORGEN DNA-kit, FADWE 96004 from BioConnect, Huissen The Netherlands), and conducted a polymerase chain reaction (PCR) (Multiplex PCR Kit from Qiagen, Hilden, Germany) to amplify five microsatellite markers selected for parental assignment (i.e. PmaC25, PmaD105, PmaGAn27, PmaTAGAn71 and PmaTGAn33; details in Saladin et al., 2003). The PCR fragments were visualised using an ABI PRISMÒ 3730 Genetic Analyser (Applied Biosystems, Foster City, CA, USA) with a molecular size standard (GeneScanTM-500 LIZÒ from Applied Biosystems, Foster City, CA, USA), and their fragment size was assessed using the software GENEMAPPER 6.0 (Applied Biosystems, Foster City, CA, USA). Chicks were assigned to be extrapair if more than one locus was not matched with the social father using an upper threshold of 3 x 10^14^ for the triolod score in CERVUS 3.0.7 (Kalinowski et al., 2007; Marshall et al., 1998). Genetic fathers were assigned to EPP chicks where possible, i.e. based on the assigned loci and spatial and temporal distance between the broods.

### Data analyses

All data processing and statistics were run in R (version 4.3.1; R Core Team, 2023) using R Studio (version 2023.06.2). To obtain diel timing of chick provisioning (i.e. after hatching), we derived the first and last detection at the nest-box entrance per day. They correspond to onsets and offsets of nest visits which were converted to onset and offset relative to sunrise and sunset, respectively. To disentangle the effect of the partner’s timing from the shared environment of the breeding pair, we ran bivariate models (*MCMCglmm* package from Hadfield, 2010) using male and female timing as response variables with Gaussian error distributions. We ran the models separately for onset (n = 634 individual-days) and offset (n = 634 individual-days), jointly referred to as *Timing. MCMCglmm* models were run with high repetition (*nitt* = 13 x 10^6^, *thin* = 10^4^, *burnin* = 3 x 10^6^) and uninformative priors (i.e. inverse-Gamma: variances set to 1, covariance to 0 and *nu* to 1.002; de Villemereuil, 2012). From the simulated data we extracted posterior means and their credible intervals (= effect sizes, *HDVinterval* function). Estimates were considered significant when their 95%-credible intervals were not overlapping with zero or with each other in regard to potential sex differences.

#### 1) Variation partitioning

We ran a bivariate *null model* to explore the magnitude of between- and within-individual variation. This information formed the basis to verify that the observed patterns represented consistent chronotypes, and that the environmental effects were truly within-individual responses. The model comprised of two separate intercepts, i.e. one for male and one for female (*Sex*), and included Pair (n = 37 of which 7 partners were not recorded) as a sex-specific random factor (*(1*|*Sex:Pair)*) to assess between-pair differences (e.g. R_Pair_). Additionally, we included Date (i.e. a categorical combination of year, month and day) as a sex-specific random factor to understand how much of the within-individual variation is explained by environmental factors. Residuals (*res*) were also modelled for both sexes separately.

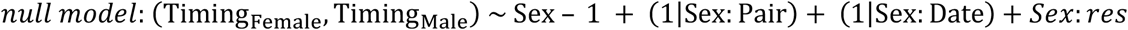

The variation partitioning was calculated for both sexes separately by dividing the variation (*σ*^*2*^) of one component (i.e. random effect or residuals) by the sum of all components (here, adjusted repeatability (*R*), Nakagawa & Schielzeth, 2010; for *MCMCglmm* see Holtmann & Dingemanse, 2022 and Houslay & Wilson, 2017). Repeatability was assessed for all random effects and the residuals.

The sex-specific R_Pair_ represented the female’s or male’s between-individual variation within pairs, e.g. comparing one female of a pair with the female from another pair. Note, that there were only six individuals measured with two different partners (i.e. in two pairs) so that R_Pair_ is a good approximation for between-individual differences in general. In contrast, within-individual variation is represented by the sum of the Date and residual repeatability.

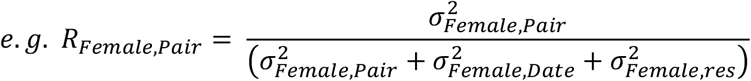

#### 2) Environmental factors

We added environmental factors that have been shown to affect diel timing previously as fixed effects to the *null model*, separately for the male and female intercepts (e.g. *Sex:Season*). In specific, we included Season (as continuous count of days from April 1^st^ = 1, i.e. April day) to look at shifts across the course of the breeding season, and ambient temperature and rainfall to assess effects of different weather conditions (Schlicht & Kempenaers, 2020). Weather data was extracted from the local weather stations (temperature from Vlieland, mean rainfall from de Kooy and Hoorn; Royal Netherlands Meterological Institute (KNMI), 2023). For ambient temperature, we used the mean temperature of the previous night (sunset to sunrise) for onset and the mean temperature of the day (sunrise to sunset) for offset. Similarly, we calculated the sum of rainfall for the previous night-time and the daytime.

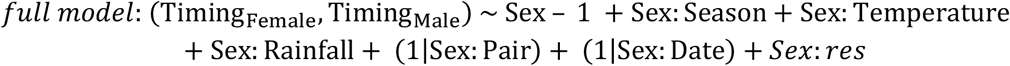

The comparison of the repeatabilities between the *null* and the *full model* can be used to investigate if the environmental effects are caused by between-or within-individual differences. However, a change in the repeatability has to be interpreted with caution because the repeatability represents a shift in the ratio of one component in relation to the total variation of random effects and residuals (see above; Nakagawa & Schielzeth, 2010; Wilson, 2018). In fact, the included fixed effects explain a part of the total variation and, therefore, they decrease the variation explained by the random factors and residuals. A stable ratio means that the variation explained by one random effect decreased proportionally with the overall variation. Likewise, an increase in repeatability represents a weaker decrease in the random effect compared to the total variation, and *vice versa*. For example, an increased individual repeatability likely shows that the environmental factors explain a part of the within-individual variation seen in the *null model*. In contrast, a reduction of the proportion of between-individual variation would indicate that individuals were only observed under different environmental factors.

In contrast to the abiotic environmental effects, we cannot extract the effect of the social partner directly from the model output. Therefore, we calculated the correlation of male and female timing (*cor*_*Pair*_) as well as the effect size (= slope) of the partner effect from the variations (*σ*^*2*^_*Female*_, *σ*^*2*^_*Male*_) and covariation (*σ*_*Female*,*Male*_) of the random factor Pair (i.e. from *full models* using the subset of the data: 29 pairs where 254 onsets and 253 offsets were measured on the same day for both pair members; Holtmann & Dingemanse, 2022 and Houslay & Wilson, 2017).

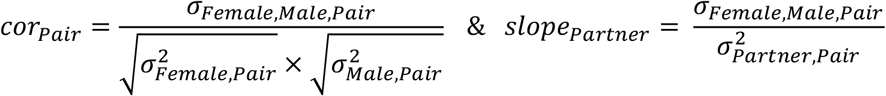

#### 3) Reproductive output

We calculated the individual’s chronotype per brood to assess the effect of male and female timing as well as their matching on the reproductive output of that brood. We used univariate generalised linear mixed effect models (*lme4* package from Bates et al., 2015) from which credible intervals were extracted based on 2,000 simulations (*bsim* function from *arm* package; Gelman & Su, 2022). For this, we mean-centred the individuals’ onsets per date to calculate diel onsets relative to conspecifics, and used for each year the individual mean of these measurements as chronotype (similar see, Schlicht et al., 2023; Steinmeyer et al., 2013; Strauß et al., pending revision.). This standardisation enabled us to compare breeding pairs measured under different environmental conditions.

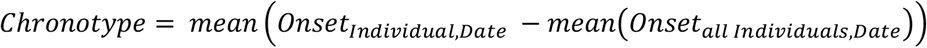

Chronotype was then used as explanatory variable together with the year-centred Hatch day, Year (as factor) and the random effect Pair. As response (*Reproductive Trait*) we used three traits in separate analyses: a) the number of fledglings (alive at chick age 14-16 days, excluding deserted broods) with a Gaussian error, b) the presence/absence of EPP chicks in the own nest using a binomial error, and c) the presence/absence of EPP chicks in other nests using a binomial error. We ran six separate models for the three response variables, and male and female chronotypes.

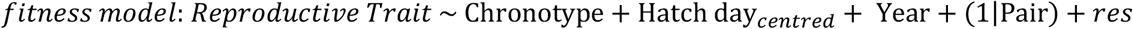

To assess the effects of chronotype matching within the breeding pair, we replaced chronotype from the previous models by the synchronisation, i.e. the absolute deviation between male and female chronotype for the corresponding brood.

## Results

### Variation partitioning of diel timing

Individuals were first detected at the nest-box entrance 14 to 20 min after sunrise, with females (13.85, 95% CI = 6.93 to 19.25) being non-significantly earlier than males (20.08, 95% CI = 14.47 to 26.37; Table S1). The opposite, and again non-significant, pattern was found for the last visit where males were detected 62 min before sunset (-62.49, 95% CI = -69.50 to -55.52) while females were about 12 min later (-50.05, 95% CI = -57.18 to -42.43). From the *null models*, we extracted between- and within-individual variation. Female onset had a repeatability of 0.37 (95% CI = 0.24 to 0.50) and male onset of 0.23 (95% CI = 0.12 to 0.35) within pairs. This was similar for offset where between-pair variation explained 0.38 (95% CI = 0.25 to 0.53) of the variation in females and 0.33 (95% CI = 0.17 to 0.49) in males. The leftover within-pair variation (0.62 to 0.77) was largely due to differences in date for onset (R_female,Date_ = 0.37, 95% CI = 0.24 to 0.49; R_Male,Date_ = 0.48, 95% CI = 0.33 to 0.61) but not as much for offset (R_Female,Date_ = 0.08, 95% CI = 0.01 to 0.16; R_Male,Date_ = 0.12, 95% CI = 0.02 to 0.22). In turn, residual variance was much larger in offset (R_Female,residual_ = 0.54, 95% CI = 0.40 to 0.66; R_Male,residual_ = 0.55, 95% CI = 0.40 to 0.70) compared to onset (R_Female,residual_ = 0.25, 95% CI = 0.18 to 0.35; R_Male,residual_ = 0.29, 95% CI = 0.20 to 0.40; for absolute variation see Table S1).

### Environmental factors

We found that individuals responded differently to environmental factors in their onset and offset of nest visits (Figure 1, Table S2). In particular, individuals significantly advanced their onset by 1 to 2 min per day relative to sunrise with the progressing breeding season. This shift was significantly stronger in females (-1.54, 95% CI = -1.90 to -1.17) than in males (-0.55, 95% CI = -1.18 to -0.04).

**Figure 1.**
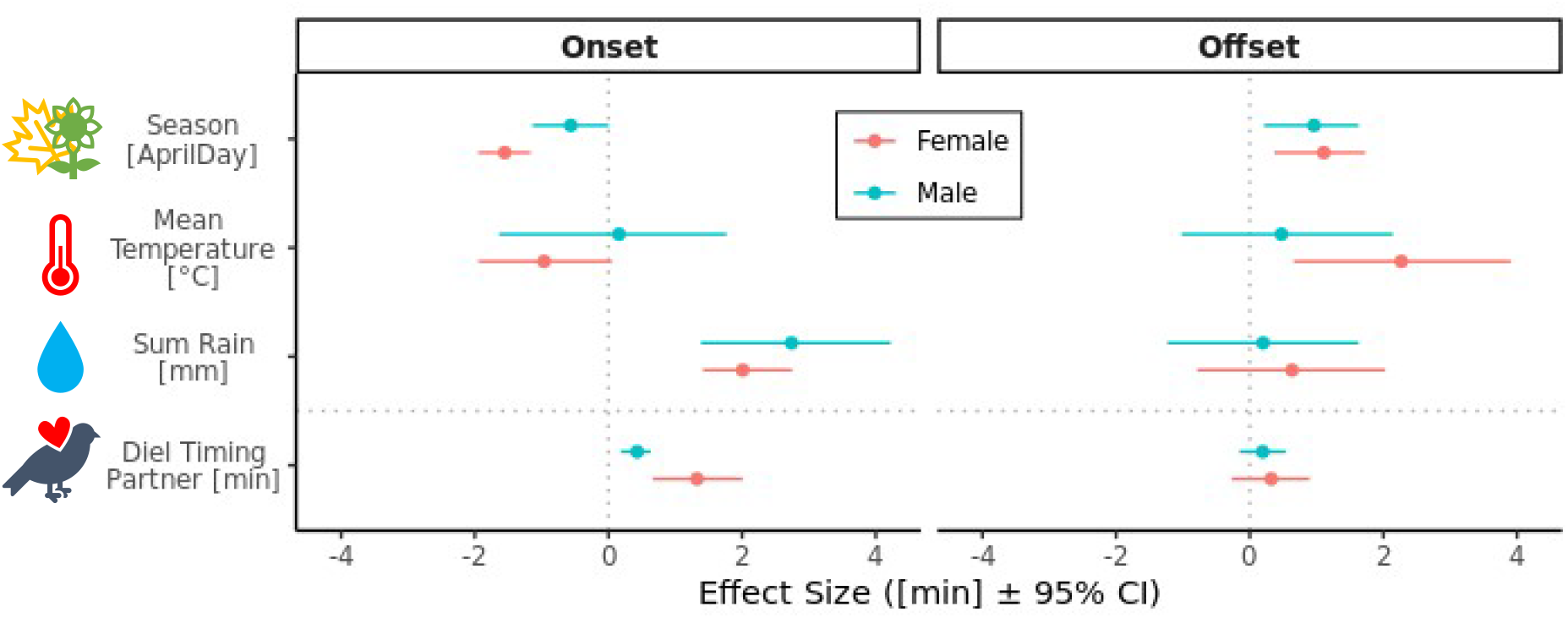
Comparison of environmental effects on onset and offset of nest visits relative to sunrise and sunset, respectively. Effect sizes (posterior means and credible intervals from the Bayesian models) are given for abiotic effects of season (with April day = days since the 31^st^ of March), ambient temperature and rainfall for the whole dataset, as well as for the effect of the partner’s timing for the subset. We used mean temperature and cumulated rainfall at night for onset and during daytime for offset.

Additionally, both sexes delayed their offset by 1 min per day relative to sunset with progressing season (female: 1.09, 95% CI = 0.39 to 1.80; male: 0.95, 95% CI = 0.29 to 1.69). While male timing did not respond to ambient temperature (onset: 0.09, 95% CI = -1.57 to 1.83; offset: 0.48, 95% CI = -1.1 to 2.18), females delayed their offset by 2 min per °C in response to warmer daytime temperatures (2.26, 95% CI = 0.71 to 3.78). Females also advanced their onset by 1 min per °C of warmer night temperature, though non-significantly (-0.99 95% CI = -1.86 to 0.11). As response to rainfall, both sexes delayed their onset significantly by 2 to 3 min per mm rainfall at night (female: 2.03, 95% CI = 1.43 to 2.72; male: 2.76, 95% CI = 1.45 to 4.23), while offset was unaffected by the rainfall during daytime (female: 0.65, 95% CI = -0.71 to 2.06; male: 0.24, 95% CI = -1.05 to 1.76). Further, we found substantial changes in the variation partitioning of female onsets when accounting for environmental conditions (for absolute variation see Table S2). The adjusted repeatability of females increased to 0.73 (95% CI = 0.61 to 0.84) within pairs and the proportional variation of date decreased accordingly to 0.07 (95% CI = 0.02 to 0.12). This reflected a corresponding increase in absolute individual variation, and a decrease in absolute variation of date (Table S1 vs. S2), indicating that females respond plastically to environmental factors rather than being measured under different environmental conditions. Similar results were obtained using a subset of the data (Table S3). Additionally, we investigated the effect of the diel timing of the partner and found a significant and high correlation of 0.74 (95% CI = 0.47 to 0.94) between female and male onsets of pairs but only a weak and non-significant one in their offsets (cor_Pair_ = 0.25, 95% CI = -0.18 to 0.62). In accordance with this, individuals shifted their onset in response to their partner’s onset. Effect size of the male response (0.43, 95% CI = 0.19 to 0.63) was significantly smaller than the one of female response (1.33, 95% CI = 0.67 to 2.01).

### Reproductive output

Investigating effects on reproductive output of the broods (Figure 2, Table S4), we found that the number of fledglings (i.e. chicks alive at age 14-16) was independent from parental chronotype (female: -0.08, 95% CI = -0.16 to 0.00; male: -0.04, 95% CI = -0.13 to 0.04) and their synchronisation (-0.01, 95% CI = -0.13 to 0.12). Female chronotype was also unrelated to the presence or absence of EPP chicks in her own nest (-0.03, 95% CI = -0.20 to 0.14) as well as to the EPP of their social male in other nests (0.00, 95% CI = -0.12 to 0.12). Similarly, there was no effect of male chronotype either (EPP in own nest: 0.04, 95% CI = -0.11 to 0.19; EPP in other nests: -0.15, 95% CI = -0.37 to 0.06). Also, the synchronisation of the parental chronotype did not explain any variation in EPP levels (EPP in own nest: -0.15, 95% CI = -13.03 to 12.80; EPP in other nests: -0.07, 95% CI = -0.31 to 0.16). We checked 109 chicks of 22 broods for EPP, and found twelve EPP chicks (11%) in six broods (29%) of which ten could be assigned to their genetic father (i.e. to 5 known extra-pair fathers). All but one EPP were observed in 2020 resulting in 19% of EPP chicks and 42% EEP broods in 2020 compared to much lower EPP levels in 2021 (2% chicks and 8% broods).

**Figure 2.**
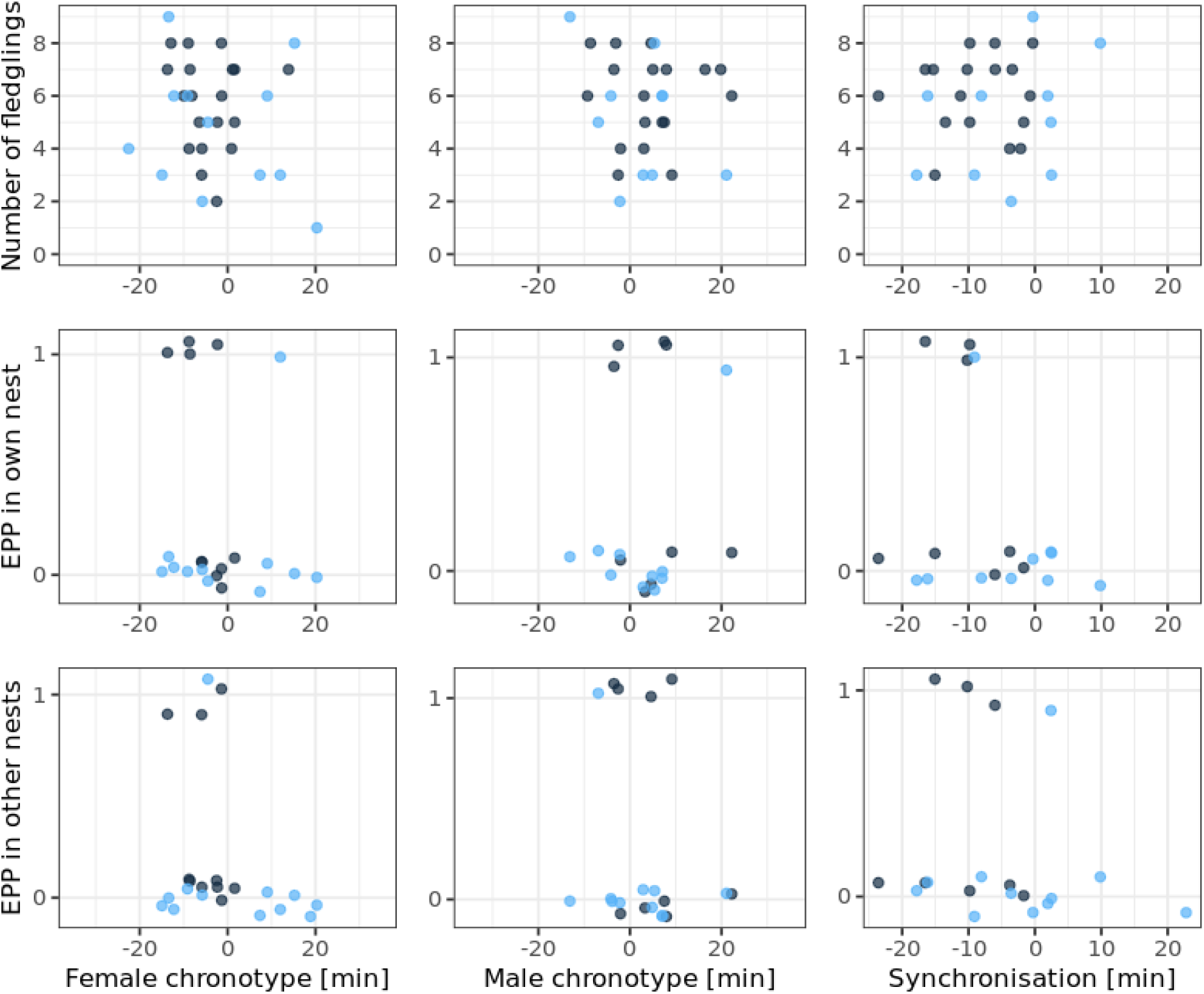
Relationship of reproductive output to parental chronotypes or within-pair synchronisation (i.e. deviation between male and female chronotype). The number of fledglings corresponds to the number of chicks alive at chick age 14-16 day. Extrapair paternity (EPP) was binomial with presence (1) or absence (0) of an EPP chick in a brood. Colours indicate the year (dark blue = 2020, light blue = 2021).

## Discussion

To explore between- and within-individual variation of diel timing, we used the onset and offset of nest visits, and found that individuals consistently differed in both traits. Further, individuals plastically advanced their onset with progressing season, and delayed it in response to rainfall and their partner’s onset, probably to adjust their timing to the environmental conditions. In contrast, individuals were less responsive to the environmental factors in their offset. The offset only delayed with the progressing season in both sexes and additionally with higher daytime temperatures in females. Other sex differences were only observed in the strength of the response to seasonal and partner effects in onset, where females responded more strongly than males. Despite the high synchronisation of the breeding pair’s onset, this synchronisation had no effect on reproductive output or EPP levels. We also could not find differences in the number of fledglings or EPP levels associated with chronotypes in males or females.

Consistent with the literature (e.g. Lehmann et al., 2012; Stuber et al., 2015) we found that individuals showed consistent differences within pairs explaining 23 to 38% of the total variation observed. By including environmental factors, the repeatability for individual female onsets increased, while the proportion of within-individual variation simultaneously decreased indicating that the observed environmental effects likely represented within-individual responses to the environment (for details see Data analyses). The effects of various environmental factors on diel timing have been investigated previously for great tits (Lehmann et al., 2012; Stuber et al., 2015) as well as for the closely related blue tits (Schlicht & Kempenaers, 2020; Steinmeyer et al., 2010). In our study, we found that individuals advanced their onset and delayed their offset of nest visits relative to sunrise and sunset, respectively, with the progressing season during chick provisioning. The lengthening of the active day could partially reflect the increase of daylength across the breeding season, though we already accounted for this by using relative onset and offset. In fact, the shift would have been even stronger using raw timing data (i.e. relative to midnight). Additionally, the birds might have responded to meet the increasing food demands of the growing chicks throughout provisioning (Barba et al., 2009; Verhulst & Tinbergen, 1997). Further, this change in activity patterns was generally less pronounced in males mostly due to the smaller advance in onset compared to the shift in females. Females shift their activity patterns already during incubation when they delay their activity onset, and often shorten their active day thereby increasing their presence on the nest (McGlade et al., 2023; Schlicht & Kempenaers, 2020; Strauß et al., 2024). This incubation shift is reversed after the hatching of the chicks and probably adds to the general season effect in females.

In response to the weather conditions, we found a delayed onset of nest visits in response to an increased amount of rain at night. This was similar to what has been previously found in blue tits (Schlicht & Kempenaers, 2020), though we did not find a response to daytime rain in offset. As a rainy night is likely associated with a rainy morning, those conditions might be unsuitable for foraging or chick provisioning. Accordingly, great tits may adjust their provisioning behaviour to avoid rainy conditions (Lejeune et al., 2019; Radford et al., 2001). In response to ambient temperature, we found no significant shift in onset as previously shown for great tits (Lehmann et al., 2012; Stuber et al., 2015) but contrasting to an advance seen in blue tits (Schlicht & Kempenaers, 2020). According to the literature, individuals’ activity offsets can shift with ambient temperature by either advancing (as observed in the lab Lehmann et al., 2012) or delaying with warmer temperatures (as observed in the wild Schlicht & Kempenaers, 2020; Stuber et al., 2015). We found a delay in offset with warmer temperatures but only in females. This sex difference in our study could be explained by sex-specific responsiveness and by a sex difference related to the methodology used for measuring activity. In particular, we used nest visits as activity measurement. Females, in contrast to males, generally roost inside the breeding nest-box (Kluijver, 1950). Therefore, the nest visits likely represent activity patterns including different types of behaviour such as foraging etc in females, while they might only reflect chick provisioning in males.

Breeding pairs synchronised their onsets of nest visits in our study. This is consistent with the two previous studies on wintering blue tits (Steinmeyer et al., 2013) and provisioning great tits (Pagani-Núñez & Senar, 2016). Steinmeyer et al. (2013) correlated the partners’ onsets measured at different timepoints before the breeding season by standardising their awakening times within date and sex. However, the positive correlation could not be replicated with the raw data from a small subset of simultaneously measured onsets (Steinmeyer et al., 2013). In contrast, Pagani-Núñez & Senar (2016) found a positive relationship using the raw onset of provisioning in great tits. This, however, comes with the problem that both parents shared the environment potentially causing this correlation (Holtmann & Dingemanse, 2022). As solution we used bivariate models to correct for the shared environment and confirmed that this correlation in onsets persists. By contrast, we found no correlation or partner effects for offset.

Individuals might respond to their partner’s onset of provisioning to match their parental effort. For example, in regard to alternation of nest visits, individuals sometimes wait for their partner to take turn in providing food to the offspring (Johnstone et al., 2014). Thus, individuals might synchronise their onset of provisioning to avoid that the partner stops and waits with provisioning in the early morning and to show reliability to the partner. Synchronised provisioning visits can be beneficial for the breeding success (Bebbington & Hatchwell, 2016; Johnstone et al., 2014; Lejeune et al., 2019; Leniowski & Wȩgrzyn, 2018; van Rooij & Griffith, 2013). Alternatively, desynchronised blue tit parents might maximise their day period for parental care resulting in higher fledgling success (Steinmeyer et al., 2013). However, we could find neither an effect of synchronised nor desynchronised onsets of the nest visits on the number of fledglings. While we only looked at one measure of fitness, parental synchronisation was also unrelated to chick body mass in blue tits (Steinmeyer et al., 2013). It is possible that the feeding rates throughout the day might be more important for fledging success. Especially the early morning provisioning seems crucial to refill energy reserves of the chicks after a night (Kontogiannis, 1967) which might be reflected in a higher number of fledglings for early females (Womack et al., 2023).

In our study there was no difference in number of fledglings between early and late individuals, neither in males nor females, which corresponds with previous findings for this population with chronotype based on incubation activity of females (Strauß et al., pending revision.). Similarly, chick condition was independent of the parental timing across various studies (Pagani-Núñez & Senar, 2016; Strauß et al., pending revision.; Womack et al., 2023). Assuming that our chronotype measurement reflects consistent timing throughout the breeding season, we expected that earliness, lateness and the synchronisation of the breeding pair was already present during the female’s fertile period. During the mating period, males are thought to anticipate the females’ timing in the morning as they arrive at the nest-box to pick up their females (Kluijver, 1950). While this seems more anecdotal, we found that, despite their overall earlier onset, females shifted three times more strongly in response to a shift in their partner’s onset than males (i.e. ca 1.5 vs. 0.5 min shift) indicating that females wait for their partner to leave the nest-box. Accordingly, both males and females have been shown to seek proximity in the early morning hours (Mace, 1989). If males were artificially delayed and missed the opportunity to guard their partner, they lose paternity in their own nest (Greives et al., 2015). In turn, synchronised diel timing might decrease EPP levels e.g. through assortative mating (Hau et al., 2017). Assortative mating is the direct or indirect selection of a partner with a similar phenotype in contrast to the adjustment towards the partner’s phenotype (Holtmann & Dingemanse, 2022). Though our sample size was unsuitable to check for assortative mating specifically (Holtmann & Dingemanse, 2022), we found no relationship between synchronisation and the EPP levels. In accordance with the literature on blue tits, we also found no support for an effect of female chronotype on extrapair paternity (Schlicht et al., 2014; Steinmeyer et al., 2013). For males, however, earliness has been found to be positively related to the number of EPP chicks in other nests (Poesel et al., 2006; Schlicht et al., 2023) which was not supported in our data (see also Steinmeyer et al., 2013). Despite a potential lack of power in our sample size, our EPP levels varied greatly between the years but were generally similar to other great tit populations with 25-30% of EPP broods and 6-7% EPP chicks (Bókony et al., 2017; Griffith et al., 2002; Roth et al., 2019; van Oers et al., 2008 but see high levels in Greives et al., 2015).

Further it is possible that also within-individual plasticity plays a crucial role for the breeding success of the pair. Plastic adjustments to the environment are adaptive when these shifts increase fitness and when individuals are exposed to changing conditions throughout their lifetime (Snell-Rood, 2013). In our study, individuals might benefit from adjusting their timing for example to the chicks’ demands, thereby increasing breeding success, and by avoiding inefficient foraging conditions during bad weather. Plasticity might be constrained by the ability of the clock to adjust to environmental (Helm et al., 2017) and other intrinsic conditions (Murren et al., 2015). This, in turn, opens up the possibility that individuals differ in their ability to be plastic (Dingemanse & Wolf, 2013). In this case, there could be even chronotype-specific plasticity in diel timing and corresponding fitness consequences, which would be interesting to investigate further in future. Overall, we could confirm a strong pair synchronisation of the onsets although we could not find any related fitness consequences for fledgling number and EPP levels.

## Conclusion

In conclusion, we observed repeatable individual differences and within-individual shifts depending on changes in the environment with a strong association of the onsets within pairs. Despite the consistent individual difference across environmental conditions, there were no differences in reproductive output. Similarly, synchronisation had no fitness consequences. However, it is possible that plasticity played an important role in maximising fitness. Adaptive plasticity enables individuals to adjust to unpredictable environmental conditions (Snell-Rood, 2013). It is not clear yet if plasticity in timing differs between individuals or even more specifically between chronotypes due to constraints by the internal clock. This might be crucial information to predict the ability for adaptation to more challenging environmental conditions (Häfker & Tessmar-Raible, 2020).

## Acknowledgment

We would like to thank Henri Bouwmeester for help with the field work, Martijn van der Sluijs for supervision of the lab work and the paternity analyses, and Erik Postma for statistical advice. We are also thankful to Staatsbosbeheer Vlieland for permission to conduct this study on their grounds. AFTS was supported by the Adaptive Life programme of the University of Groningen, the Netherlands.

BMT was supported by a NWO VENI grant (VI.Veni.192.022) and by the The Arctic Seasonal Timekeeping Initiative (ASTI) grant from UiT strategic funds.

## Data Availability Statement

Data and codes are available upon request and will be deposited in a repository upon publication.

