## Supplementary Material for "Variation in diel timing in great tits is affected by the timing of their social mate"

Table S1: Variation partitioning. Model estimates and credible intervals of provisioning onset or offset in minutes relative to sunrise or sunset, respectively. Estimates were extracted from posteriors of the null models (i.e. excluding environmental factors) for fixed and random effects. 95%-Credible intervals that do not overlap with zero indicate significance. Sample sizes are given as the number of measurements per sex.

| **Onset** | | **Female** n = 360 | | | | | **Male** n = 274 | | | |
| --- | --- | --- | --- | --- | --- | --- | --- | --- | --- | --- |
| **Fixed effects** | | **β^2^** | | **95% CI** | | | **β^2^** | | **95% CI** | |
| Intercept | | 13.85 | | (6.93, 19.25) | | | 20.08 | | (14.47, 26.37) | |
| **Random** | **Variation Female** | | | | **Variation Male** | | | **Covariation** | | |
| **effects** | **σ^2^** | | **95% CI** | | **σ^2^** | **95% CI** | | **σ_xy_** | | **95% CI** |
| Pair | 196.63 | | (105.37, 315.33) | | 118.35 | (46.34, 197.20) | | 126.02 | | (48.83, 216.85) |
| Date | 193.73 | | (108.39, 287.83) | | 244.85 | (134.79, 365.92) | | 175.97 | | (99.30, 282.27) |
| Residual | 128.49 | | (104.66, 150.80) | | 146.84 | (113.56, 174.83) | | 31.22 | | (1.00, 59.05) |
| **Offset** | | **Female** n = 363 | | | | | **Male** n = 271 | | | |
| **Fixed effects** | | **β^2^** | | **95% CI** | | | **β^2^** | | **95% CI** | |
| Intercept | | -50.05 | | (-57.18, -42.43) | | | -62.49 | | (-69.50, -55.52) | |
| **Random** | **Variation Female** | | | | **Variation Male** | | | **Covariation** | | |
| **effects** | **σ^2^** | | **95% CI** | | **σ^2^** | **95% CI** | | **σ_xy_** | | **95% CI** |
| Pair | 445.23 | | (225.79, 726.24) | | 303.97 | (114.57, 524.91) | | 108.54 | | (-87.92, 315.29) |
| Date | 95.37 | | (13.31, 196.69) | | 109.54 | (17.21, 214.25) | | 96.59 | | (15.17, 177.97) |
| Residual | 607.36 | | (506.70, 714.63) | | 495.24 | (381.52, 602.64) | | 203.92 | | (123.47, 315.96) |

Table S2: Environmental factors. Model estimates and credible intervals of provisioning onset or offset in minutes relative to sunrise or sunset, respectively, for the full models including environmental factors. We used mean temperature and cumulated rainfall at night-time. Estimates were extracted from the posteriors for fixed and random effects. 95%-Credible intervals that do not overlap with zero indicate significance. Sample sizes are given as the number of onsets or offsets per sex.

| **Onset** | | | **Female** n = 360 | | | **Male** n = 274 | | | |
| --- | --- | --- | --- | --- | --- | --- | --- | --- | --- |
| **Fixed effects** | | | **β^2^** | **95% CI** | | **β^2^** | | **95% CI** | |
| Intercept | | | 115.82 | (94.71, 137.63) | | 49.85 | | (16.15, 83.24) | |
| Season [April day] | | | -1.54 | (-1.90, -1.17) | | -0.55 | | (-1.18, -0.04) | |
| Night Temperature [°C] | | | -0.99 | (-1.86, 0.11) | | 0.09 | | (-1.57, 1.83) | |
| Night Rainfall [mm] | | | 2.03 | (1.43, 2.72) | | 2.76 | | (1.45, 4.23) | |
| **Random** | **Variation Female** | | | **Variation Male** | | | **Covariation** | | |
| **effects** | **σ^2^** | **95% CI** | | **σ^2^** | **95% CI** | | **σ_xy_** | | **95% CI** |
| Pair | 370.35 | (199.69, 568.04) | | 116.68 | (44.64, 196.20) | | 170.57 | | (73.53, 284.91) |
| Date | 34.04 | (13.86, 53.91) | | 153.20 | (80.14, 236.80) | | 68.80 | | (35.61, 104.42) |
| Residual | 117.31 | (98.55, 135.86) | | 148.15 | (119.70, 178.37) | | 31.73 | | (8.58, 59.91) |
| **Offset** | | | **Female** n = 363 | | | **Male** n =271 | | | |
| **Fixed effects** | | | **β^2^** | **95% CI** | | **β^2^** | | **95% CI** | |
| Intercept | | | -146.48 | (-187.49, -109.36) | | -125.87 | | (-166.74, -87.12) | |
| Season [April day] | | | 1.09 | (0.39, 1.80) | | 0.95 | | (0.29, 1.69) | |
| Day Temperature [°C] | | | 2.26 | (0.71, 3.78) | | 0.48 | | (-1.10, 2.18) | |
| Day Rainfall [mm] | | | 0.65 | (-0.71, 2.06) | | 0.24 | | (-1.05, 1.76) | |
| **Random** | **Variation Female** | | | **Variation Male** | | | **Covariation** | | |
| **effects** | **σ^2^** | **95% CI** | | **σ^2^** | **95% CI** | | **σ_xy_** | | **95% CI** |
| Pair | 556.21 | (269.37, 883.71) | | 271.59 | (99.08, 474.19) | | 125.37 | | (-105.14, 307.45) |
| Date | 62.83 | (0.83, 128.35) | | 74.93 | (0.36, 154.67) | | 63.13 | | (-1.51, 123.53) |
| Residual | 563.92 | (466.86, 657.08) | | 517.50 | (406.99, 636.29) | | 220.14 | | (122.09, 320.14) |

##### Table S3: Environmental factors. Model estimates and credible intervals from the full models using a subset of the data where both partners were measured simultaneously at the same day. Estimates were extracted from the posteriors for fixed and random effects. 95%-Credible intervals that do not overlap with zero indicate significance. Sample sizes are given as the number of onset and offsets.

| **Onset** n = 254 | | | **Female** | | | **Male** | | | |
| --- | --- | --- | --- | --- | --- | --- | --- | --- | --- |
| **Fixed effects** | | | **β^2^** | **95% CI** | | **β^2^** | | **95% CI** | |
| Intercept | | | 89.94 | (67.92, 113.30) | | 30.11 | | (-1.25, 64.34) | |
| Season [April day] | | | -1.12 | (-1.50, -0.71) | | -0.27 | | (-0.92, 0.28) | |
| Night Temperature [°C] | | | -0.78 | (-1.73, 0.07) | | 0.39 | | (-1.37, 2.22) | |
| Night Rainfall [mm] | | | 1.85 | (1.12, 2.59) | | 2.55 | | (1.15, 3.90) | |
| **Random** | **Variation Female** | | | **Variation Male** | | | **Covariation** | | |
| **effects** | **σ^2^** | **95% CI** | | **σ^2^** | **95% CI** | | **σ_xy_** | | **95% CI** |
| Pair | 276.23 | (127.16, 467) | | 90.73 | (34.09, 163.300) | | 118.35 | | (31.97, 227.90) |
| Date | 28.20 | (11.35, 46.03) | | 161.05 | (89.07, 241.99) | | 62.84 | | (32.39, 97.19) |
| Residual | 69.04 | (54.14, 83.20) | | 126.53 | (99.44, 155.10) | | 16.20 | | (2.60, 31.70) |
| **Offset** n =253 | | | **Female** | | | **Male** | | | |
| **Fixed effects** | | | **β^2^** | **95% CI** | | **β^2^** | | **95% CI** | |
| Intercept | | | -143.69 | (-184.57, -103.87) | | -101.55 | | (-141.74, -59.88) | |
| Season [April day] | | | 0.85 | (0.13, 1.54) | | 0.57 | | (-0.19, 1.34) | |
| Day Temperature [°C] | | | 3.22 | (1.56, 4.72) | | 0.33 | | (-1.36, 2.10) | |
| Day Rainfall [mm] | | | 0.70 | (-0.67, 2.13) | | -0.13 | | (-1.71, 1.26) | |
| **Random** | **Variation Female** | | | **Variation Male** | | | **Covariation** | | |
| **effects** | **σ^2^** | **95% CI** | | **σ^2^** | **95% CI** | | **σ_xy_** | | **95% CI** |
| Pair | 523.30 | (251.63, 916) | | 317.50 | (124.39, 531.60) | | 101.60 | | (-78.04, 313.60) |
| Date | 69.39 | (7.40, 137.7) | | 74.46 | (0.45, 142.00) | | 67.74 | | (9.48, 125.10) |
| Residual | 357.90 | (286.38, 439.40) | | 432.10 | (340.56, 525.60) | | 120.50 | | (61.83, 194.00) |

##### Table S4: Reproductive output. Estimates for nine reproductive models studying a) the number of fledglings (i.e. chick alive at age 14-16 days), b) the presence (1) or absence of extrapair paternity (EPP) in the own nest or c) the presence (1) or absence of male EPP other nests, in relation to female and male chronotype and their synchronisation. The model estimates and credible intervals were calculated by bootstrapping for fixed (β^2^) and random effects (σ^2^). Residual variance was fixed to 1 in binomial models for presence/absence of EPP. Credible intervals that do not overlap with zero indicate significance. Sample sizes for every model are given as the number of broods.

| **a) Number of fledglings** | **Female** n = 31 | | **Male** n = 28 | | **Synchronisation** n = 27 | |
| --- | --- | --- | --- | --- | --- | --- |
|  | **β^2^** | **95% CI** | **β^2^** | **95% CI** | **β^2^** | **95% CI** |
| Intercept_2020_ | 5.378 | (4.430, 6.326) | 6.006 | (5.014, 7.018) | 6.070 | (4.548, 7.539) |
| Chronotype | -0.075 | (-0.157, 0.004) | -0.043 | (-0.129, 0.037) | -0.007 | (-0.132, 0.117) |
| Hatch day | 0.134 | (0.021, 0.249) | 0.063 | (-0.050, 0.175) | 0.064 | (-0.052, 0.179) |
| Year_2021_ | -1.049 | (-2.509, 0.456) | -0.994 | (-2.604, 0.592) | -1.107 | (-2.789, 0.504) |
|  | **σ^2^** | **95% CI** | **σ^2^** | **95% CI** | **R** | **95% CI** |
| Pair | 0.000 | (0.000, 0.000) | 0.000 | (0.000, 0.000) | 0.000 | (0.000, 0.000) |
| Residual | 3.857 | (2.256, 6.675) | 3.955 | (2.180, 7.222) | 3.889 | (2.160, 7.014) |
| **b) EPP in**  **own nest** | **Female** n = 21 | | **Male** n = 19 | | **Synchronisation** n = 18 | |
|  | **β^2^** | **95% CI** | **β^2^** | **95% CI** | **β^2^** | **95% CI** |
| Intercept_2020_ | -0.504 | (-2.021, 1.117) | -0.468 | (-2.07, 1.183) | -9.161 | (-182.992, 165.179) |
| Chronotype | -0.033 | (-0.203, 0.140) | 0.041 | (-0.110, 0.185) | -0.145 | (-13.028, 12.797) |
| Hatch day | 0.071 | (-0.133, 0.280) | 0.037 | (-0.157, 0.236) | 0.038 | (-14.527, 14.853) |
| Year_2021_ | -2.153 | (-4.994, 0.675) | -2.290 | (-5.262, 0.667) | -1.007 | (-177.639, 181.706) |
|  | **σ^2^** | **95% CI** | **σ^2^** | **95% CI** | **σ^2^** | **95% CI** |
| Pair | 0.364 | (0.166, 0.645) | 0.751 | (0.345, 1.325) | 1615.200 | (673.186, 2968.208) |
| **c) EPP in**  **other nests** | **Female** n = 23 | | **Male** n = 20 | | **Synchronisation** n = 19 | |
|  | **β^2^** | **95% CI** | **β^2^** | **95% CI** | **β^2^** | **95% CI** |
| Intercept_2020_ | -1.259 | (-3.015, 0.571) | 0.372 | (-1.479, 2.178) | -0.292 | (-3.959, 3.344) |
| Chronotype | -0.001 | (-0.119, 0.117) | -0.149 | (-0.369, 0.060) | -0.072 | (-0.306, 0.164) |
| Hatch day | -0.146 | (-0.445, 0.156) | -0.107 | (-0.364, 0.146) | -0.211 | (-0.618, 0.195) |
| Year_2021_ | -1.293 | (-3.982, 1.405) | -2.965 | (-6.304, 0.407) | -1.657 | (-4.999, 1.514) |
|  | **σ^2^** | **95% CI** | **σ^2^** | **95% CI** | **σ^2^** | **95% CI** |
| Pair | 0.058 | (0.029, 0.098) | 0.000 | (0.000, 0.000) | 0.414 | (0.191, 0.719) |
